# Visual impairment cell non-autonomously dysregulates brain-wide proteostasis

**DOI:** 10.1101/2023.10.19.563166

**Authors:** Shashank Shekhar, Katherine J Wert, Helmut Krämer

## Abstract

Loss of hearing or vision has been identified as a significant risk factor for dementia but underlying molecular mechanisms are unknown. In different Drosophila models of blindness, we observe non-autonomous induction of stress granules in the brain and their reversal upon restoration of vision. Stress granules include cytosolic condensates of p62, ATF4 and XRP1. This cytosolic restraint of the ATF4 and XRP1 transcription factors dampens expression of their downstream targets during cellular stress. Cytosolic condensates of p62 and ATF4 were also evident in the thalamus and hippocampus of mouse models of congenital or degenerative blindness. These data indicate conservation of the link between loss of sensory input and dysregulation of stress responses critical for protein quality control in the brain.

**One-sentence summary:** Drosophila and mouse models link loss of visual input to dysregulated stress responses of neurons and glia in the brain.

## Main text

In addition to genetic causes, potentially modifiable factors significantly contribute to the risk for dementia (*1, 2*). For example, hearing loss in midlife is estimated to contribute to 8.2% of world-wide dementia cases (*1*) and vision impairment is associated with 1.8% of US dementia cases (*2*). Together these studies highlight the importance of maintaining sensory input during ageing but the mechanistic link between the loss of sensory input and the onset or progression of dementia remains unknown.

A common aspect of different neurodegenerative disorders, including dementia, is dysregulated protein homeostasis or proteostasis (*3–7*). In response to different cellular stressors, proteostasis is maintained by a network of interdependent protein quality control mechanisms (*8*), including the ubiquitin proteasomal system, autophagy and the integrated stress response (ISR) (*9*). Proteostasis is particularly crucial in brain health because neurons need to be maintained for an entire lifespan, they have intricate morphology which defines their participation in complex neural circuits, and the dynamic regulation of pre and postsynaptic sites imposes localized constrains on their protein turnover (*10, 11*). A prominent stress response is the assembly of neuronal stress granules (SGs). SGs are membrane-less liquid-liquid phase-separated ribonucleoprotein organelles in the cytoplasm (*12, 13*). SGs can serve a transient protective function (*14*), followed by their dissolution once homeostasis is restored (*15, 16*). Under chronic stress, however, persistent SGs accumulate additional markers such as p62 (also known as Sequestosome-1 in mammals or Ref(2)p in Drosophila) and may serve as seeds for the aggregation of proteins related to neurodegenerative diseases (*13*). Here, we explore the dysregulation of proteostasis as molecular link between blindness and neurodegeneration.

## Results

### Disruption of photoreceptor synaptic function causes brain-wide proteostasis dysregulation

We previously (*17*) observed disrupted proteostasis in the visual system upon expression of dominant-negative Shibire (Shi^ts1^) or Tetanus Toxin Light chain (TTL) under control of a photoreceptor-specific driver (Fig. 1A). Unexpectedly, the resulting inhibition of synaptic activity specifically in photoreceptors triggered proteostasis defects throughout the brain. This was visualized by a staining for ATF4 and XRP1, two transcription factors induced by the ISR (*18*). For both, we detected densely stained punctae in cell bodies throughout the brain, in numbers significantly increased compared to wild-type controls (Fig. 1B-F). Furthermore, condensates of Ref(2)p/p62, a recently appreciated marker for proteostasis defects (*19*), were also significantly increased throughout the brain of flies with blocked photoreceptor cell synapses but not wild-type controls (Fig. 1B’’-D’’,G).

**Fig. 1.**
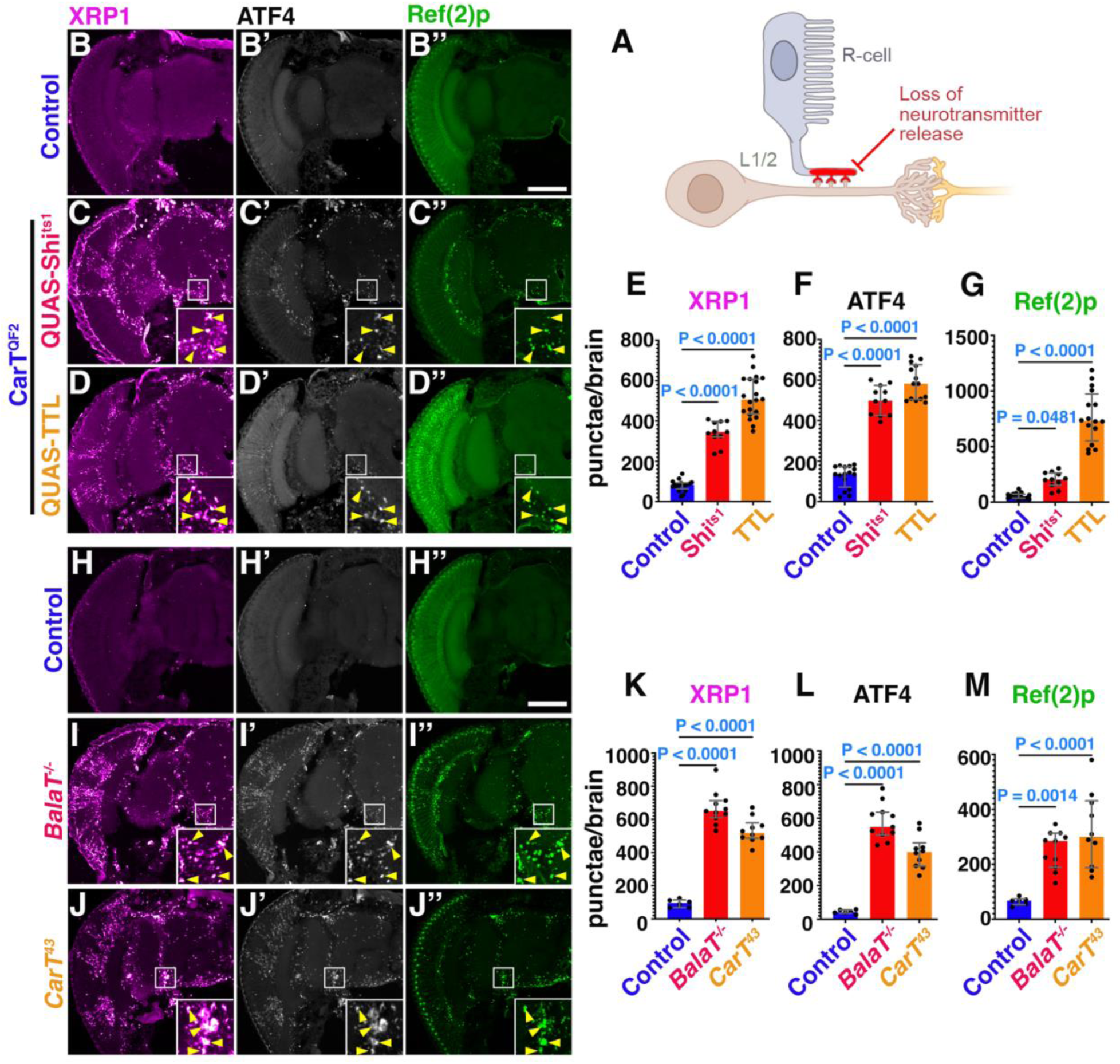
Loss of photoreceptor synaptic activity causes formation of condensates of p62, ATF4 and XRP1 throughout the brain. (A) Schematic representation of approach used in figure 1 to disrupt photoreceptor activity. (B-B’’) Micrographs of cryosections of adult brain control (QUAS-Shi^ts^), immunostaining for XRP1 transcription factor (B), ATF4 (B’) and Ref(2)p, also known as Drosophila p62 (B’’), quantified in (E-G). Insets are 3x magnified from the indicated area and adjusted for brightness and contrast to better visualize condensates. Yellow arrows indicate condensates which colocalize. (C-C’’) Representative adult head cryosections of flies expressing Shi^ts1^ in photoreceptors. Staining for XRP1 is shown in (C), ATF4 in (C’), and Ref(2)p condensates in (C’’) with respective quantification in E,F, and G. (D-D’’) Cryosections of adult fly heads expressing TTL in photoreceptors to block synaptic release. Immunostaining for XRP1 (D), ATF4 (D’) and Ref(2)p (D’’) is quantified in E,F, and G. (H-J’’) Micrographs of cryosections of adult brains from controls (QUAS-Shi^ts^) (H), or of mutants inhibiting histamine neurotransmitter recycling in *BalaT* (I) and *CarT* (J) heads immunostained for XRP1 (I), ATF4 (I’), and Ref(2)p condensates (I’’) quantified in K, L and M. Bar graphs show median with interquartile range and data repented are pooled from 3 independent sets of experiments. Significance threshold was determined by one-way ANOVA with Bonferroni correction for multiple comparisons. P-values are represented in each graph. Scale bar in B-D’’ and H-J’’ is 100 µm. Genotypes are listed in Suppl. Table 1.

To further test non-autonomous effects of blocked retinal synaptic activity, we interfered with recycling of histamine, the neurotransmitter of photoreceptor neurons. The CarT and BalaT transporters are required in photoreceptor neurons or glia, respectively, for histamine neurotransmitter recycling and the corresponding mutant animals are functionally blind (*20–23*). In these animals, we found significant brain-wide increases of ATF4, XRP1 and Ref(2)p punctae in both transporter mutants compared to wild-type controls (Fig. 1H-M).

Together, our data demonstrate that disruption of photoreceptor synaptic output dysregulates proteostasis in cells deep in the brain.

### Restored vision rescues proteostasis in neurons and glia of blind *norpA* flies

To further test the hypothesis that blindness non-autonomously dysregulates brain proteostasis, we used *norpA* mutants (Fig. 2A). The *norpA* gene encodes Phospholipase-c beta which catalyzes an early step in the phototransduction pathway (*24*); *norpA* mutants have been extensively used to study functional consequences of blindness in flies (*25*). In *norpA* brains, punctae of ATF4, XRP1 and p62 were prevalent (Fig. 2B,D-F). As part of canonical ISR activation, elevated levels of the ATF4 and XRP1 enter the nucleus and drive a transcriptional program that aids in restoring cellular homeostasis (*18*). By contrast, in brains of blind flies, ATF4 and XRP1 accumulate in punctae (Fig. 2G) that resemble cytosolic stress granules (SGs), typically assembled by liquid-liquid phase separation of ribonucleoproteins (*13*). Within these granules, ATF4 closely colocalized with XRP1 and Ref(2)p and was not associated with the nucleus (Fig. 2G,H). Consistent with the presence of XRP1 in SGs of *norpA* brains, we observed co-localization with the RNA-binding proteins Caprin and dFMR1 (Fig. 2I-K), two markers of SGs (*26, 27*). Surprisingly, XRP1/ATF4-positive granules in *norpA* brains were negative for Rin (suppl Fig. 1 A-H), the Drosophila ortholog of G3BP which is commonly associated with SGs (*27–29*). In *norpA* brains, XRP1-positive SGs were present in neurons, marked by elav staining and in glia marked by repo-Gal4 driven mCD8::RFP (suppl Fig. 1H-K).

**Fig. 2.**
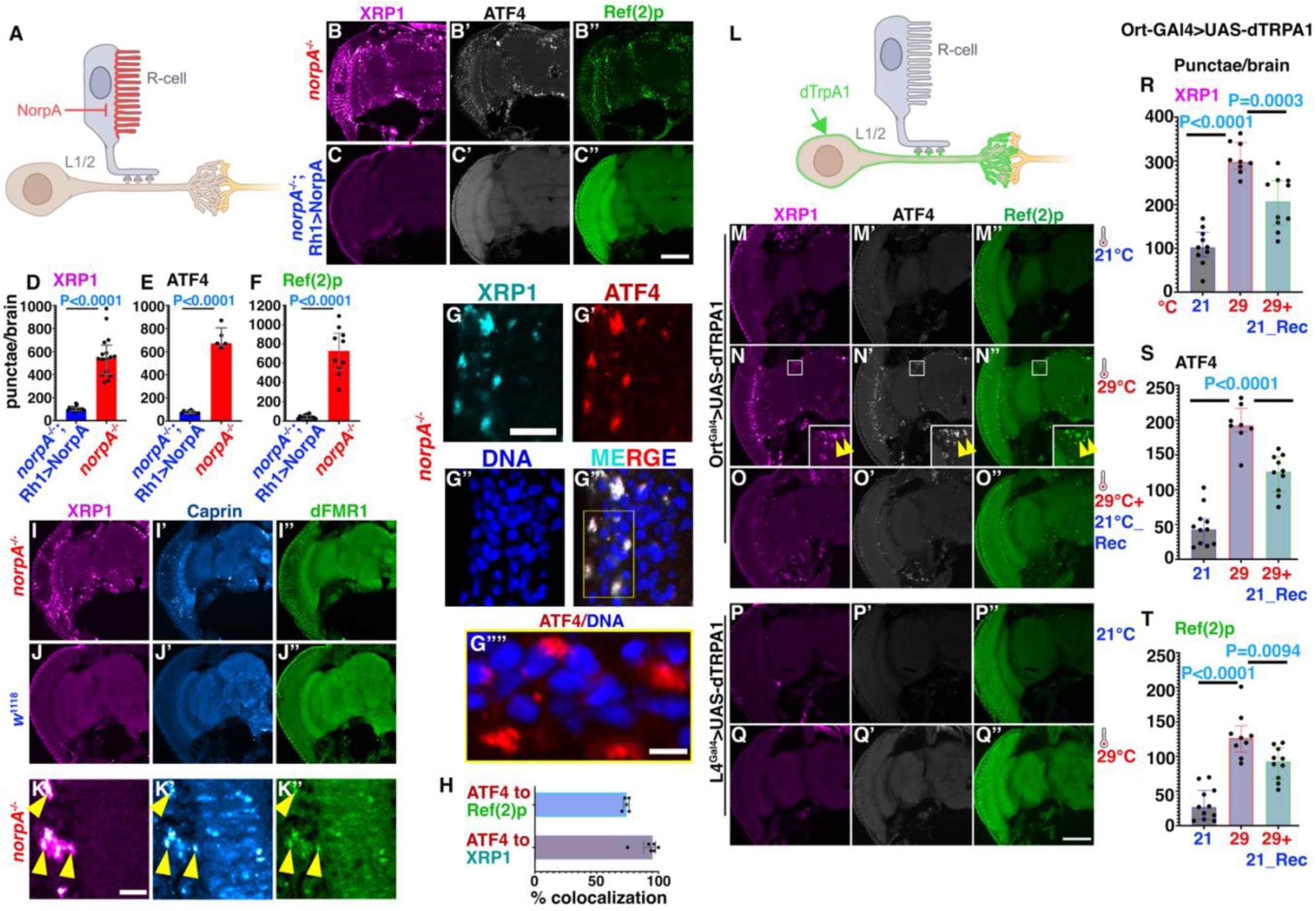
Visual impairment triggers stress granule formation non-autonomously in the brain. (A) Schematic representation of *norpA* based approach to disrupt photoreceptor activity in (B-H). (B-B’’) Representative image of adult *norpA* head cryosections (B). XRP1 condensates, quantified in (D), ATF4 (B’), quantified in (E) and Ref(2)p condensates (B’’), quantified in (F). (C-C’’) Eye-specific rescue of *norpA* by Rh1 promoter-driven expression. XRP1(C), ATF4 (C’) and Ref(2)p (C’’), quantified in (D), (E), and (F), respectively. Bar graphs show median with interquartile range of number of punctae per fly brain. Data are pooled from 3 independent experiments. Statistical significance threshold was determined by Two-tailed Unpaired t test and P-Value is represented in each graph. (G-H) Colocalization and cytosolic restraint of XRP1 and ATF4 in *norpA* brains. XRP1 (G) and ATF4 (G’) colocalize in cytosolic condensates outside of nuclei marked by DNA (G’’,G’’’). Yellow box in (G’’’) is magnified in (G’’’’) to show cytosolic location of ATF4. (H) Quantification of colocalization of ATF4 with XRP1 and ATF4 with Ref(2)p. (I-K) Cryosections of *norpA* or *w*^1118^ control brains stained for XRP1 (I-K), Caprin (I’-K’) and dFMR1 (I’’-K’’). Representative images shown from two independent experiments. (K) High-magnification images show partial colocalization as indicated by yellow arrowheads. (L) Schematic representation of UAS-dTRPA1 expression-based approach to hyperactivate lamina neurons and disrupt visual circuits in (M-T). (M-Q) Micrographs of adult hemi-brains expressing UAS-dTRPA1 under control of ort-Gal4 (M-O) or L4-Gal4 (P,Q) drivers maintained at 21°C (M-M”, P-P”), activated at 29°C (N’-N”, Q-Q”) or recovered from 29°C at 21°C (O--O’’). Staining for XRP1, (M-Q,R) ATF4 (M’-Q’,S) and Ref(2)p (M’’-Q’’,T) revealed reversible induction of brain-wide condensates by ort-Gal4 driven dTRPA1 expression at elevated temperature, but not by L4-Gal4 driven expression. Functional expression for both was verified in Suppl Fig. 2. Bar graphs (R-T) show median with interquartile range of number of punctae. Data are pooled from 2 independent sets of experiments. Significance threshold was determined by one-way ANOVA with Bonferroni correction for multiple comparisons and P-Value is represented in each graph. Scale bar in B-C’’, G-H’’ and M-Q’’ is 100 µm, I-J’’’ is 10 µm. Genotypes are listed in Suppl. Table 1

Importantly, the presence of ATF4, XRP1 and Ref(2)p punctae was effectively suppressed (Fig. 2C,D-F) when vision in *norpA* mutants was restored by photoreceptor-specific expression of NorpA under control of the Rh1 rhodopsin promoter (*30*). The number of punctae of ATF4, XRP1 and Ref(2)p in these rescued flies was reduced to near wild-type levels (Fig. 2D-F, compare to Fig. 1B,G). Together, these results indicate that blindness and the resulting lack of photoreceptor synaptic activity results in SGs in neurons and glia throughout the brain and that restoring vision prevents formation of these SGs.

### Hyperactivation of lamina neurons causes SG formation in the brain

Photoreceptor synapses release histamine as an inhibitory neurotransmitter and therefore loss of photoreceptor synaptic output may hyperactivate downstream lamina neurons in the phototransduction circuit (*31*). To mimic this situation in vivo, we used a thermogenetic approach: activation of lamina neurons by the temperature-sensitive dTRPA1 channel (Fig. 2L). To express UAS-dTRPA1 specifically in synaptic targets of photoreceptors we used the ort-Gal4 driver which is expressed in multiple classes of lamina neurons that receive direct input from photoreceptors. Activation of dTRPA1 at 29°C in these lamina neurons significantly increased brain-wide ATF4, XRP1 and p62 condensates compared to flies kept at 21°C (Fig. 2M,N,R-T). Moreover, three days after shifting these flies from 29°C back to 21°C, the number of brain-wide punctae for ATF4, XRP1 and p62 were significantly reduced (Fig. 2O,R-T) indicating that blindness-induced SGs are reversible although with a significant delay.

Unlike Ort-positive neurons, L4 lamina neurons do not receive direct input from photoreceptors (*32*). L4 hyperactivation by dTRPA1 expression at 29°C did not cause elevated SG levels in brains, which were indistinguishable from those maintained at 21°C (Fig. 2P,Q). This indicates that SG formation is not a generic effect of neuronal hyperactivation. L4-expressed dTRPA1 is functional, nevertheless, as indicated by the altered sleep behaviors triggered by the shift to 29°C for Ort- and L4-expressed dTRPA1 (suppl. Fig. 2). Together, these data suggest that blindness-induced SG formation is reversible and specific to a subset of visual circuits.

### Sequestering ATF4 and XRP1 in SGs dampens their canonical response to stress

To test whether SG-mediated sequestration of ATF4 and XRP1, two key effectors of the ISR, affects the response to cellular stress, we used a transcriptional reporter expressing LacZ under control of the Thor promoter which is responsive to ATF4 activity (*18*). To induce cellular stress, we fed flies tunicamycin (TM) at a concentration known to induce the ISR (*33*). Whereas wild-type brains displayed significant TM-induced enhancement of Thor-lacZ expression in both dorsal and ventral clusters of neuronal cell bodies (Fig. 3A-C,F), the number of Thor-lacZ expressing cells was significantly lower in TM-fed *norpA* brains (Fig. 3D-F). Similar changes were observed for GstD-GFP, a reporter for XRP1 activity (*18*). Western analysis of whole-head lysates revealed TM-induced increase of GstD-GFP expression in wild-type flies (Fig. 3G,H). Compared to wild-type flies exposed to vehicle-control only, corresponding *norpA* flies show elevated level of GFP but failed to further increase GFP expression upon TM feeding (Fig. 3G,H). In sections of TM-fed *norpA* brains, cells with prominent XRP1-positive SGs exhibited negligible activation of the GstD-GFP reporter, in contrast to cells lacking SGs (indicated with dashed lines in Suppl Fig. 3A-F). Despite the elevated ATF4 and XRP1 staining in SGs in *norpA* brains, their mRNA levels were not significantly different from controls (Suppl Fig. 3G). Furthermore, downstream transcriptional targets of the ISR trended lower (Suppl Fig. 3G). To test whether this dysregulation of the ISR may be linked to neurodegeneration, we measured the number of brain vacuoles in two-week old brains (*34*). Compared to photoreceptor-rescued *norpA*;Rh1>NorpA flies, *norpA* mutants displayed elevated levels of brain vacuoles (Fig. 3I-K). Altogether, these data suggest that sequestration of ATF4 and XRP1 in stress granules compromises their canonical transcriptional function and dysregulated stress responses in *norpA* flies are linked to elevated neurodegeneration.

**Fig. 3.**
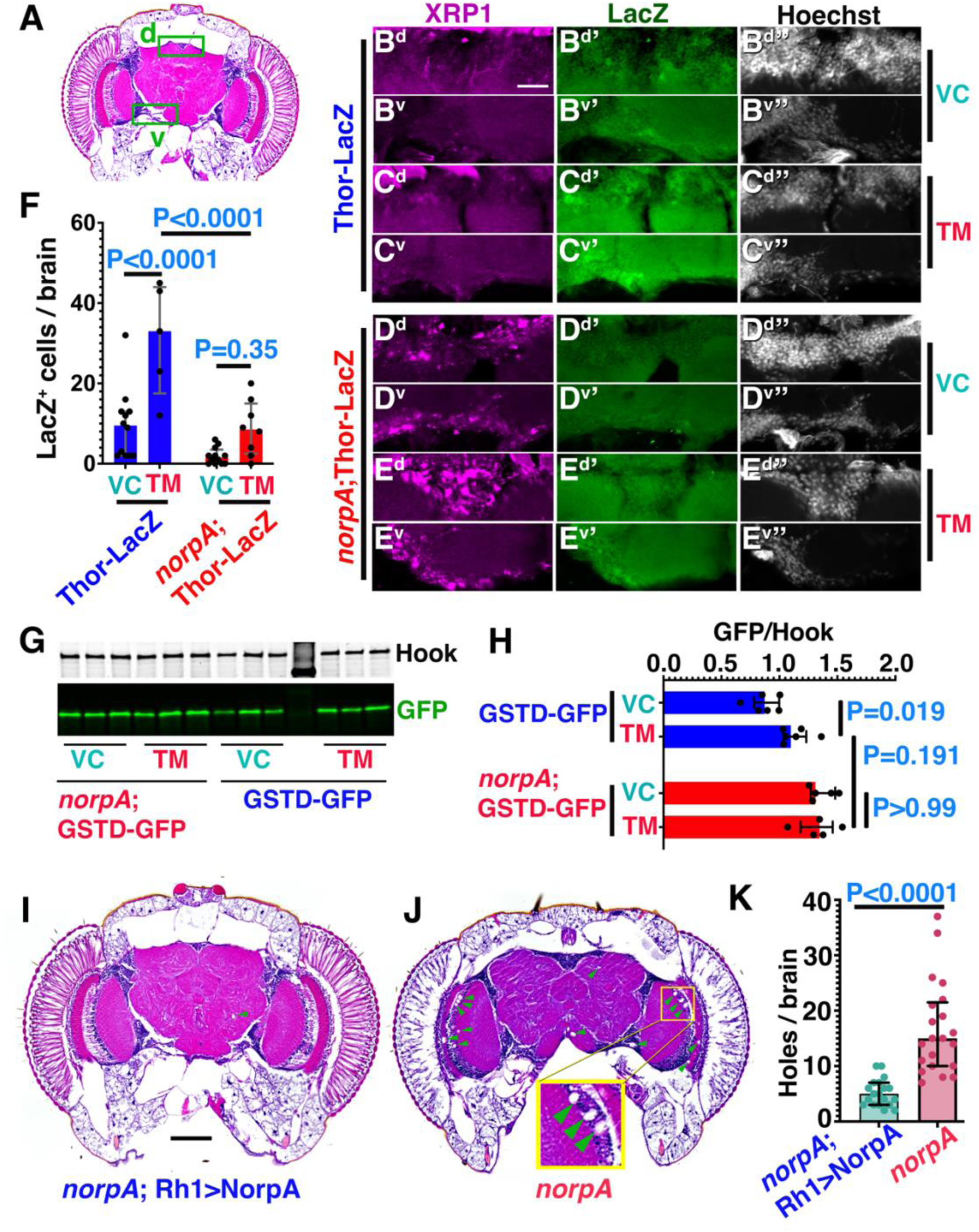
Sequestration of ATF4 and XRP1 in stress granules dampens their stress responsiveness. (A) Schematic of fly brain pointing to dorsal (d) and ventral (v) regions imaged in panels (B-E). (B-E) Adult brain cryosections of Thor-lacZ (B,C) or *norpA*;Thor-lacZ (D,E) brains from flies fed DMSO (0.02%) as vehicle control (VC) or tunicamycin (TM, 12 µM) as indicated and stained for lacZ, XRP1 and DNA. *S*cale bar in B^d^ is 25 µm in C and F. (G-H) Representative western blot (G) with three biological repeats per treatment condition of either 12 µM tunicamycin (TM) or 0.02% DMSO as vehicle control (VC). Hook serves as loading control. Quantification of blots is represented in (H), bar graphs show median with interquartile range of fold change of GFP signal upon TM feeding to adult flies. P-values from two-way ANOVA followed by Bonferroni correction for multiple comparisons test are represented in the bar graphs. (I-K) H&E-stained brain sections of two-week old *norpA*; Rh1>NorpA (I) and *norpA* (J) flies were assessed for brain vacuoles indicative of neurodegeneration with quantification in (K). Inset in (J) shows magnified examples of brain vacuoles, Scale bar in I is 50 µm. Genotypes are listed in Suppl. Table 1.

### Blindness induced non-autonomous proteostasis dysregulation in brain is conserved in mice

The visual systems of *Drosophila* and mammals share conserved design principles and mechanisms of synaptogenesis and neuronal circuit formation (*35*). We wondered whether this similarity extends to the blindness-induced dysregulation of proteostasis in the brain. In the thalamus and hippocampus of congenital blind *Vsx2* mice (*36, 37*) we observe increased condensates of p62 (Fig. 4A-D,E,I) and significantly elevated ATF4 levels (Fig. 4F-H), especially in the cytosolic staining of ATF4 (Fig. 4H’’) when compared to wild-type 129S1/SvlmJ controls (Fig. 4G’’).

**Fig. 4.**
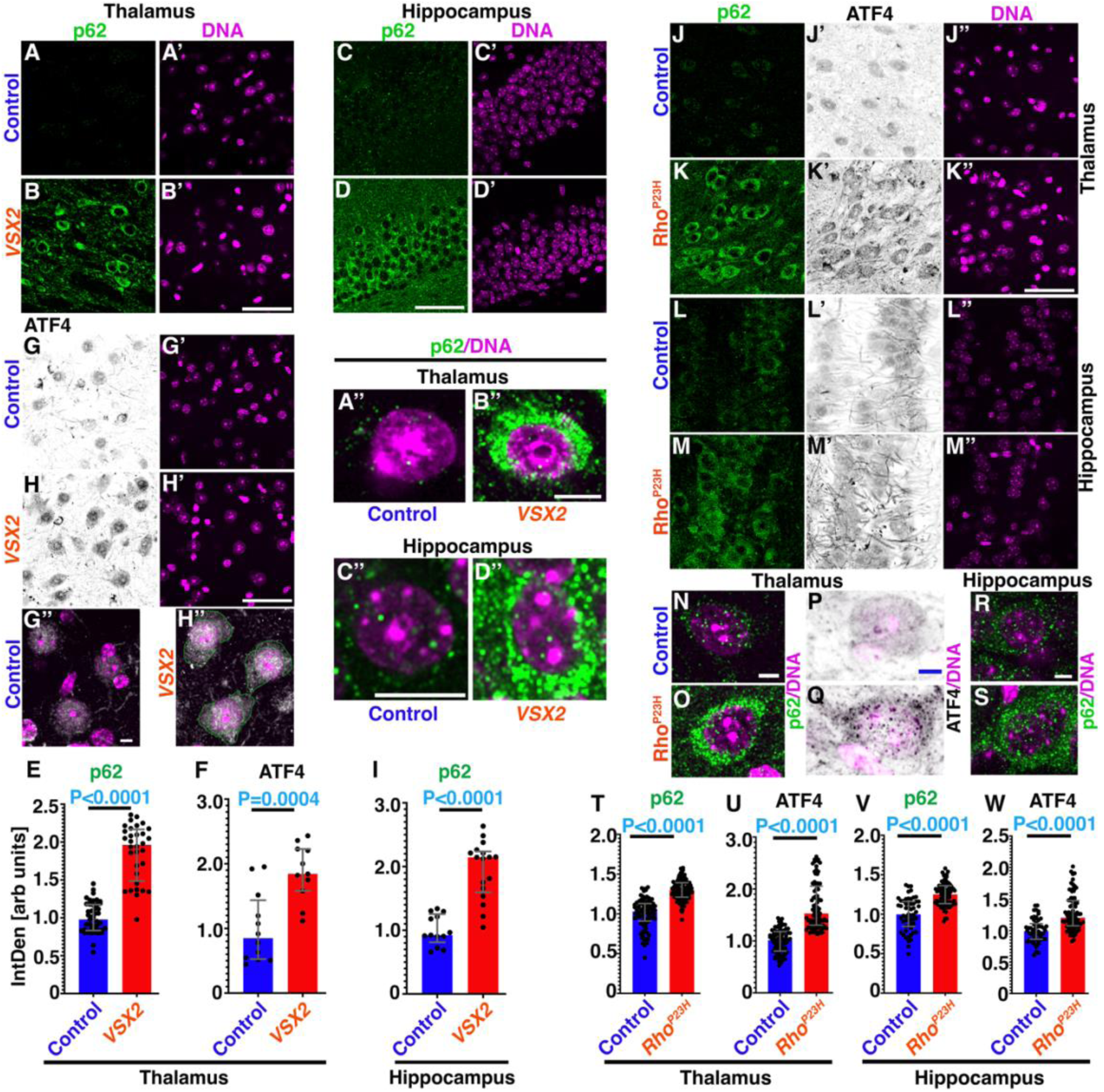
Specific visual circuit neuron in lamina regulates SGs formation in brain and visual impairment induced SGs brain-wide is evolutionary conserved in mice. (A-I) Section of thalamus (A,B,G,H; quantified in E,F) and hippocampus (C,D; quantified in I)) of adult, congenitally blind *Vsx2* mice (B,D,H) and age-matched 129S1/SvlmJ controls (A,C,G) stained for p62 (A-D) or ATF4 (G-H). Magnified, merged images highlight increased cytosolic staining for p62 (A’’-D’’) and ATF4 (G’’,H’’) in *Vsx2* brains. (H’’) Dotted green line indicates cytosolic content. Bar graphs (E-F, I) show median with interquartile range of integrated density. Data are pooled from 4 independent biological replicates. Significance threshold was determined by Two-tailed Unpaired t test and P-values shown in each graph. (J-S) Sections from 10-week-old B6/J control (J,L) and degenerative *Rho*^P23H^ (K,M) brains stained for p62 (J-M) and ATF4 (J’-M’). Regions shown are thalamus (J,K, quantified in T,U) and hippocampus (L,M, quantified in V,W). (N-S) Elevated cytosolic staining is revealed in magnified, merged images showing p62/DNA (N,O) or ATF4/DNA (P,Q) in thalamus and p62/DNA (R,S) in hippocampus. Bar graphs (T-W) show median with interquartile range of integrated density. Data are pooled from 4 independent biological replicates. Significance threshold was determined by Two-tailed Unpaired t test and P-values shown in each graph. Scale bars in B’, D, H’ and K’’ is 50 µm for A-D, G-H and J-M. Scale bars in G”, N and R is 5 µm in G,H, N-S. Scale bar in B’’ is 8 µm in A’’,B’’. Scale bar in C’’ is 10 µm in C’’,D’’. Genotypes are listed in Supplementary Table 1.

To test whether similar consequences arise from retinal degeneration, we used *Rho*^P23H^ mice that model the most common rhodopsin mutant in humans linked to retinal degeneration (*38*). In 10-week-old homozygous *Rho*^P23H^ mice, we detected increased levels of cytosolic ATF4 and p62 condensates in thalamus and hippocampus when compared to C57BL/6J (B6/J) wildtype controls (Fig. 4J-W, suppl Fig. 4).

Together, these data indicate a link between visual impairment and dysregulated brain proteostasis that is conserved from flies to mammals.

## Discussion

Once neurons are stressed, activation of the ISR as well as the formation of stress granules are both double-edged swords. ISR-mediated eIF2α phosphorylation lightens the load of unfolded proteins by reducing general translation and elevates ATF4 levels thereby promoting expression of a range of transcripts that aid in cellular stress adaptation (*9, 39*). Upon persistent activation, however, the induced expression of CHOP in mammals and XRP1 in Drosophila complicates the outcomes, as their transcriptional targets not only promote stress recovery but also can induce cell death (*18, 39–43*). Similarly, SGs can serve a transient protective function by limiting translation of a subset of mRNAs and confining caspases, (*12, 14, 28*), but their persistent presence can also promote neurodegeneration (*13*).

In blind flies, we observe an unexpected merger of these two stress responses with the two ISR-inducible transcription factors ATF4 and XRP1 being highly enriched within SGs. This confinement counteracts their elevated levels as we observe reduced expression of ATF4 and XRP1 target genes in blind flies, especially in cells with prominent ATF4/XRP1-positive SGs. Such restrained ISR activation is probably beneficial in the absence of exogenous stressors. However, limited induction under stress conditions, as we observe in TM-fed flies, constitutes a vulnerability. Indeed, links between dysregulated ISR and neurodegenerative diseases are well established (*3, 9, 44, 45*).

Consistent with this notion, dysregulated proteostasis is a hallmark of neurodegenerative diseases (*3, 5*), typically triggered by interactions with aggregation-prone proteins encoded by diseases-specific risk genes, such as Amyloid-β, Tau, α-Synuclein, Huntingtin or SOD1 (*46*). Our data in fly brains show that persistently altered circuit activity due to loss of sensory input dysregulates proteostasis thereby reducing stress resistance and contributing to the risk of dementia in ageing. Elevated cytosolic ATF4 and p62 condensates in brains of mouse models of retinal degeneration (*Rho*^P23H^) and congenital blindness (*Vsx2*) suggest an underappreciated role of the connection between the loss of sensory input and non-autonomous proteostasis regulation and the consequences for the progression of neurodegenerative diseases.

## Acknowledgments

We thank Drs. Todd Roberts, Kim Huber, Peter Douglas and members of the Krämer lab for helpful comments on the manuscript. We are grateful to Michael Ewnetu and Charles Tracy for technical assistance with sleep and qRT-PCR experiments, and Jose Cabrera for help with figures. We thank Drs. Hyung Don Ryoo and Joseph M. Bateman, and the Developmental Studies Hybridoma Bank at The University of Iowa for antibodies and the Bloomington Drosophila Stock Center (NIH P40OD018537) for flies.

## Author contributions

Conceptualization: SS, HK

Methodology: SS, HK, KJW

Investigation: SS

Funding acquisition: HK, KJW

Supervision: HK, KJW

Writing – original draft: SS

Writing – review & editing: SS, HK, KJW

## Data and materials availability

All data are available in the main text or the supplementary materials.

## Supplementary Data

### Material and Methods

#### Biological reagents

| Reagent type (species) or resource | Designation | Source or reference | Identifiers | Additional information |
| --- | --- | --- | --- | --- |
| genetic reagent (*D. melanogaster*) | QUAS-Shi<sup>ts</sup> | Bloomington Drosophila Stock Center | BDSC:30013; FBgn0003392; RRID:BDSC_30013 |  |
| genetic reagent (*D. melanogaster*) | QUAS-TTL | Bloomington Drosophila Stock Center | BDSC: 91808; FBgn0016753; RRID:BDSC_91808 |  |
| genetic reagent (*D. melanogaster*) | UAS-dTRPA1 | Bloomington Drosophila Stock Center | BDSC: 26264<br>FBti0114502<br>RRID:BDSC_26264 |  |
| genetic reagent (*D. melanogaster*) | w<sup>1118</sup> | Bloomington Drosophila Stock Center | BDSC:3605; FBti0131930; RRID:BDSC_3605 |  |
| genetic reagent (*D. melanogaster*) | Balat[D-Gal4] | this study |  |  |
| genetic reagent (*D. melanogaster*) | w[*] *norpA*[P24]; P{w+=ninaE-norpA.W}2 | Bloomington Drosophila Stock Center | BDSC: 52276; FBst0052276<br>RRID:BDSC_52276 |  |
| genetic reagent (*D. melanogaster*) | w [*] *norpA*[P24] | Bloomington Drosophila Stock Center | BDSC: 9048<br>FBst0009048<br>RRID:BDSC_9048 |  |
| genetic reagent (*D. melanogaster*) | repo-Gal4 | Bloomington Drosophila Stock Center | BDSC: 7415<br>FBst0009048<br>RRID:BDSC_7415 |  |
| genetic reagent (*D. melanogaster*) | UAS-mCD8::RFP | Bloomington Drosophila Stock Center | BDSC: 32218<br>FBst0009048<br>RRID:BDSC_32218 |  |
| genetic reagent (*D. melanogaster*) | *CarT*[43] | (22) | FBgn0032879 |  |
| genetic reagent (*D. melanogaster*) | *CarT*<sup>HA-T2A-QF2</sup> | (17) | FBgn0032879 |  |
| genetic reagent (*D. melanogaster*) | L4-Gal4 | Bloomington Drosophila Stock Center | BDSC: 49883<br>RRID:BDSC_49883 |  |
| genetic reagent (*D. melanogaster*) | Ort-Gal4 | Bloomington Drosophila Stock Center | BDSC: 56517<br>RRID:BDSC_56517 |  |
| genetic reagent (*D. melanogaster*) | Ort-Gal4 | Bloomington Drosophila Stock Center | BDSC: 56515<br>RRID:BDSC_56515 |  |
| genetic reagent (*D. melanogaster*) | GSTD-GFP | (47) | Gift from Dr. Don Ryoo, NYU |  |
| antibody | anti-GFP (Chicken polyclonal) | ThermoFisher Scientific | ThermoFisher Scientific:A10262; RRID: AB_2534023 | 1:500 IHC |
| antibody | anti-GFP (Mouse monoclonal, B2) | Santa Cruz | Santa Cruz Anti-GFP Antibody (B-2): sc-9996 | 1:250 WB |
| antibody | anti-P62/SQSTM1 (rabbit polyclonal) | Proteintech Group, Inc<br>Catalog Number: 18420-1-AP |  | 1:200 IHC |
| antibody | Anti-Ref (2)p (rabbit polyclonal) | This paper |  | 1:3000 IHC |
| antibody | Alexa 488- or 568- or 647 secondaries<br>Alexa Fluor 568 Goat anti-mouse<br>Alexa Fluor 488 Goat anti-mouse | ThermoFisher | thermoFisher fluorescent-secondary-antibodies | 1:500 IHC |
|  | Alexa Fluor 647 Goat anti-mouse<br>Alexa Fluor 488Goat anti-rabbit<br>Alexa Fluor 568 Goat anti-rabbit<br>Alexa Fluor 647 Goat anti-rabbit<br>Alexa Fluor 647 Goat anti-guinea pig<br>Alexa Fluor 568 Goat anti-guinea pig<br>Alexa Fluor 488 Goat anti-guinea pig<br>Alexa Fluor 668 Goat anti-rat<br>Alexa Fluor 488 Goat anti-chicken |  |  |  |
| antibody | IRDye 800CW Goat anti mouse<br>IRDye 700DX Goat anti mouse<br>IRDye 800CW Goat anti Rabbit<br>IRDye 700DX Goat anti Rabbit | LICOR Biosciences | https://www.licor.com/bio/reagents/new-reagent=category=irdye-secondary-antibodies | 1:20,000 WB |
| antibody | anti-RFP (Rabbit polyclonal) | Rockland | Rockland:600-401-379; RRID:AB_2209751 | 1:500 IHC |
| antibody | anti-dFMR1(Rabbit polyclonal) | Developmental Studies Hybridoma Bank | RRID: AB_528252 | 1:500 IHC |
| antibody | anti-Rin (Rabbit polyclonal) | (48) | Gift from Dr. Elizabeth R Gavis | 1:500 IHC |
| antibody | anti-Caprin (Rabbit polyclonal) | (49) | Gift from Dr.Ophelia Papoulas UT Austin, | 1:500 IHC |
| antibody | anti-XRP1 (Guinea Pig polyclonal) | (18) | Gift from Dr. Don Ryoo, NYU | 1:1000 IHC |
| antibody | anti-ATF4 (mouse monoclonal) | Proteintech Group, Inc Catalog Number: 60035-1-Ig |  | 1:500 IHC |
| antibody | Anti-Crc (ATF4 rat polyclonal) | (50) | Gift from Dr. Joseph Bateman, King's College | 1:2000 IHC |
| antibody | Anti-Repo (mouse monoclonal) | Developmental Studies Hybridoma Bank | RRID:AB_528448 | 1:500 IHC |
| antibody | Anti-ELAV (Rat monoclonal) | Developmental Studies Hybridoma Bank | RRID:AB_528217 | 1:500 IHC |
| antibody | Anti-Hook (rabbit polyclonal) | (51) | lab generated | 1:5000 WB |

#### Fly rearing conditions

Flies used in this study were reared on standard molasses fly food, under room temperature conditions. For tunicamycin treatments, 1% sucrose solution in 1.3% agarose was used at 25°C. All flies were aged 3-5 days and placed in 5 cm diameter vials containing normal food, with no more than 30 flies. Head dissections were performed between 1 PM and 4 PM. For temperature sensitive experiments, temperature used is indicted in corresponding figures. The *Balat*^Δ-Gal4^ allele was generated by replacing the *Balat* coding sequences with Gal4 using CRISPR/Cas9. Detailed analysis of the allele will be published elsewhere.

#### Mouse lines and husbandry

All experiments were performed in accordance with the ARVO Statement for the Use of Animals in Ophthalmic and Visual Research and were all approved by the Animal Care and Use Committee at University of Texas Southwestern Medical Center. B6.129S6(Cg)-*Rho^tm1.1Kpal^*/J mice, herein referred to as *Rho*^P23H^ mice, and 129S1/Sv-*Vsx2^or-J^*/J mice, herein referred to as *Vsx2* mice, were obtained from the Jackson Laboratory (Bar Harbor, ME, USA) (RRID:IMSR_JAX:017628; RRID: IMSR_JAX:000395). Age-matched C57BL/6J (B6/J) mice (RRID:IMSR_JAX:000664) or age-matched 129S1/SvlmJ mice (RRID:IMSR_JAX002448) were used as experimental controls. Mice were bred and maintained at the facilities of the University of Texas Southwestern Medical Center. Animals were kept on a light–dark cycle (12 hour–12 hour). Food and water were available *ad libitum* throughout the experiment and all brain samples were collected between the hours of 1 PM and 4 PM.

##### Immunohistochemistry

Fly heads were dissected as described previously (*17*). Briefly, after proboscis removal, heads were immediately transferred to ice-cold 3% glyoxal solution pH 4.0 (*52*) for one hour and washed overnight in 25% (wt/vol) sucrose in phosphate buffer (pH 7.4), embedded in Optimal Cutting Temperature compound (EMS, Hatfield, PA) and quickly frozen in liquid nitrogen and sectioned at 20-μm thickness on a cryostat microtome (CM 1950, Leica Microsystems, Wetzlar, Germany). Sections were washed three times (PBS with 0.1% triton X100, PBST), blocked (10% NGS) and probed with primary antibody diluted in 5% NGS solution. Antibody dilutions are listed in the key biological reagents table. Images were taken with a 20x NA-0.8 or 40X NA0.95 WD 0.17-0.25mm FOV 25mm Air Objective or an oil-immersion 63× NA-1.4 lens on an inverted confocal microscope (LSM710 or LSM880 with Airyscan, Carl Zeiss or Nikon AXR confocal). For each genotype, immunohistochemistry experiments were performed in three biological replicas with independent sets of flies, using identical acquisition settings. Experimenters were blinded to sample identity before acquiring images or before quantification as appropriate. Polyclonal antibodies against the p62 peptide PRTEDPVTTPRSTQ (*53*) were raised in rabbits and affinity purified (Genemed Synthesis, Inc).

For collection of mouse brains for immunohistochemistry, mice were anesthetized by intraperitoneal injection of 0.2 mL/10 g body weight of 1 mL of 100 mg/mL ketamine with 0.1mL of 20mg/mL xylazine diluted in 8.9mL 1X PBS. Mice were then transcardially perfused with 1X PBS, followed by 10% formalin. Brains were removed and fixed in 10% formalin at 4°C overnight. Brains were then embedded in 4% low-gelling temperature agarose (Sigma Aldrich) and 150 µm serial sections were collected into 1X PBS using the Leica VT1000S vibrating blade microtome.

For IHC of mise brain, sections were gently transferred to fresh 24 well plate and washed five time with PBST. Sections were blocked with 10%NGS for 2 hrs and probed with primary antibodies diluted in 5% NGS solution. After three PBST washes, secondary antibody incubation and three more PBST washes, sections were float-mounted on Probe On Plus microscope slides (Fisher Scientific). Antibody dilutions are listed in the key biological reagents table. Images were taken with a 20x NA-0.8 or an oil-immersion 63× NA-1.4 lens on an inverted confocal microscope (LSM880 with Airyscan, Carl Zeiss).

##### Quantification of fluorescence staining

Fluorescence images were quantified using ImageJ (NIH) adapting previous methods (*54*) and Imaris 9.5 (Oxford Instruments) software was used for punctate quantification and colocalization analysis. For each antibody, a threshold was determined, removing the lowest signal in control samples (to reduce variation from low level background signals). This same threshold was applied to a mask created for every image in a batch of staining. For the quantification of ATF4 and p62 in mouse brains, within 1-µm optical slices, regions were selected manually and assigned as Regions of Interest. The integrated pixel intensity per unit area was measured within this selected area. For the quantification of ATF4 and XRP1 in fly brain, Imaris software was used to quantify punctae in respective images in 3D by creating surface Ref(2)p was quantified by spot counting, regions were selected manually and assigned as Regions of Interest. The same Regions of Interest are used for each experimental condition. For Thor-LacZ, images were AI denoised using Nikon NIS-Elements. Denoised images were used for quantification, wild-type TM-fed images were used to set the threshold for LacZ^+^ cells, the same threshold was used to count cells in other groups.

##### H&E staining and quantification

To assess neurodegeneration, fly heads were dissected and after proboscis removal fixed in 4% PFA for 48 hrs, embedded in paraffin and 5 µm sections were cut followed by H&E staining as described (*34*). Images were taken with a 20X objective using a Nikon Fi3 color Camera (NA0.75 WD 1.0mm FOV 25mm). Following blinding for genotype and treatment, images were quantified using ImageJ; after thresholding of the ROI, vacuoles in the brain structure were counted with a cutoff of holes more than 5µm in size (*34*).

##### Western blot

For western blotting, 3-5 days aged 10 adult fly heads were homogenized in 100 µl lysis buffer (10% SDS, 6 M urea, and 50 mM Tris-HCl, pH 6.8) and then kept rotating for 30 min at 4°C followed by boiling at 95°C, for 2 min. Lysates were spun for 10 min at 15,000x*g*, supernatant was separated. Before loading on gel 1mM DTT was added in samples followed by boiling at 95°C, for 2 min, 5 µl (1/2 heads) of lysates were separated on 4-20 % gradient SDS-PAGE, transferred to nitrocellulose membrane, blocked with 3% non-fat dried milk dissolved in wash buffer (20 mM Tris-HCl pH 7.5, 5.5 mM NaCl and 0.1 % Tween-20) and probed in primary antibodies in following dilutions; rabbit anti-Hook (1:5000), Mouse anti-GFP clone B2 (1: 250), and incubated overnight at 4°C on rotator. Blots were washed 3 times 10 mins each with wash buffer. Bound antibodies were detected and quantified using IR-dye labelled secondary antibodies (1:15,000) and the Odyssey scanner (LI-COR Biosciences). Pre-stained molecular weight markers (HX Stable) were obtained from UBP-Bio.

##### Tunicamycin feeding

For inducing ER stress, fly food was prepared by adding tunicamycin (TM) at a final concentration of 12 µM. TM was added in 1% sucrose solution with1.3% low melting agarose at temperature 40°C. Flies aged 3 to 5 days were transferred to food containing either TM or DMSO as vehicle control (VC) and placed in the 12h:12h LD chambers at 25°C for 23 hrs. After completion of the treatment, flies were used for western blot or immunofluorescence.

##### Measuring sleep

Sleep was measured as described previously (*17*). Briefly, 3-5 days old male flies were placed inside a 65 x 3 mm glass tube with standard molasses fly food. Tubes were placed in the Ethoscope arena (*55*). Ethoscopes were placed in a 21°C incubator with 300 Lux light at a 12h:12h LD cycle and after allowing flies to acclimatize for one day, their movement was recorded. On the third day, the temperature was shifted to 29°C to activate the temperature-sensitive dTRPA channel. After 3 days at 29°C, the temperature was shifted back to 21°C for recovery. Ethoscope data were analyzed using rethomics (*56*). All codes used in the sleep analysis are available in github (https://github.com/mz27ethio/Ethoscope-Kramerlab.git).

##### Statistical analysis

Statistical significance of results was analyzed using Prism-GraphPad10. Anderson-Darling or Shapiro-Wilk test were used to assess the normality assumption for continuous parameters. Afterwards, skewed data were transformed on log scale followed by similar normality assessment. Independent t-test was used to compare normally distributed parameters between two groups whereas Mann-Whitney U test served as non-parametric test.

For comparing groups of three and more, one-way analysis of variance followed by Bonferroni correction (a multiple comparisons test) was used for normally distributed sample parameters. Two-way analysis of variance followed by Bonferroni’s test was used for multiple comparisons to identify the individual pairwise comparisons to separate the effects of treatment and genetic backgrounds. Skewed parameters were compared using Kruskal-Wallis test followed by Dunn’s test for multiple comparison test. Bar graphs resulted from these analyses demonstrate median with interquartile range. P values smaller than 0.05 were considered as statistically significant, and values are indicated in the corresponding graphs.

### Supplemental Figures

**Suppl Fig. 1.**
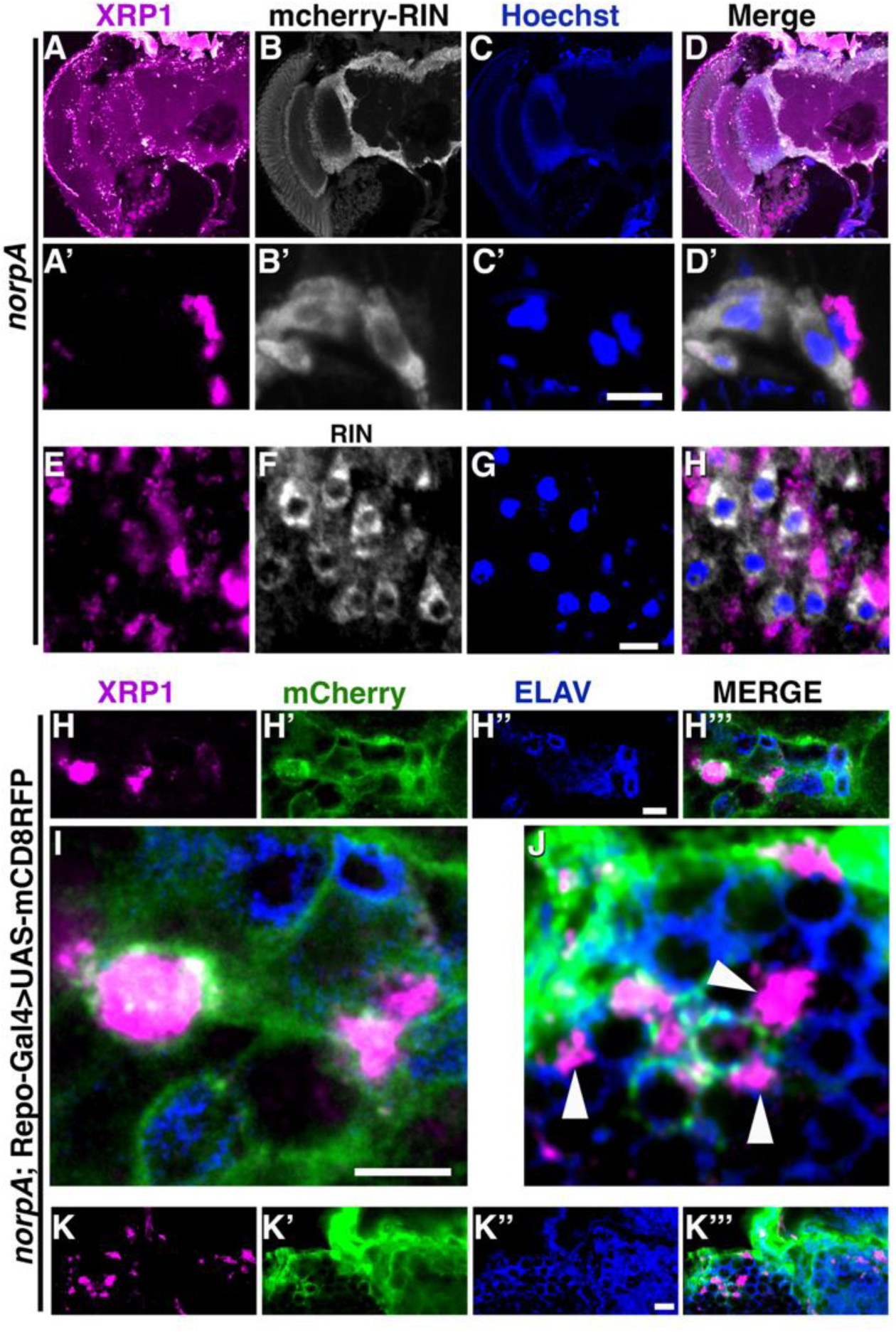
Blindness-induced SGs do not colocalize with RIN/G3BP. (A-D) Cryosections of *norpA;* mCherry-Rin fly brain immunostained for XRP1 (A), mCherry (B) and DNA (C). Merged image shown in (D). (A’-D’) High resolution details show exclusion of mCherry-Rin from XRP1-positive SGs. (E-H) Cryosections of *norpA* head immunostained for XRP1 (E), endogenous RIN protein (F) and DNA (G). Merged image in (H) shows exclusion of Rin protein from XRP1-positive SGs. (H-K) Cryosections of heads of *norpA* flies expressing UAS-mCD8-RFP under control of the glial repo-Gal4 driver stained for XRP1 (H,K), mCherry (H’,K’), and neuronal ELAV protein (H’’,K’’); merged images (H’’’,K’’’). Magnified images (I,J) show XRP1-positive SGs within glia and neurons (white arrowheads in J). Scale bars in C’, G, and H’’ are 5 µm. Genotypes are listed in Supplementary Table 1.

**Suppl Fig. 2.**
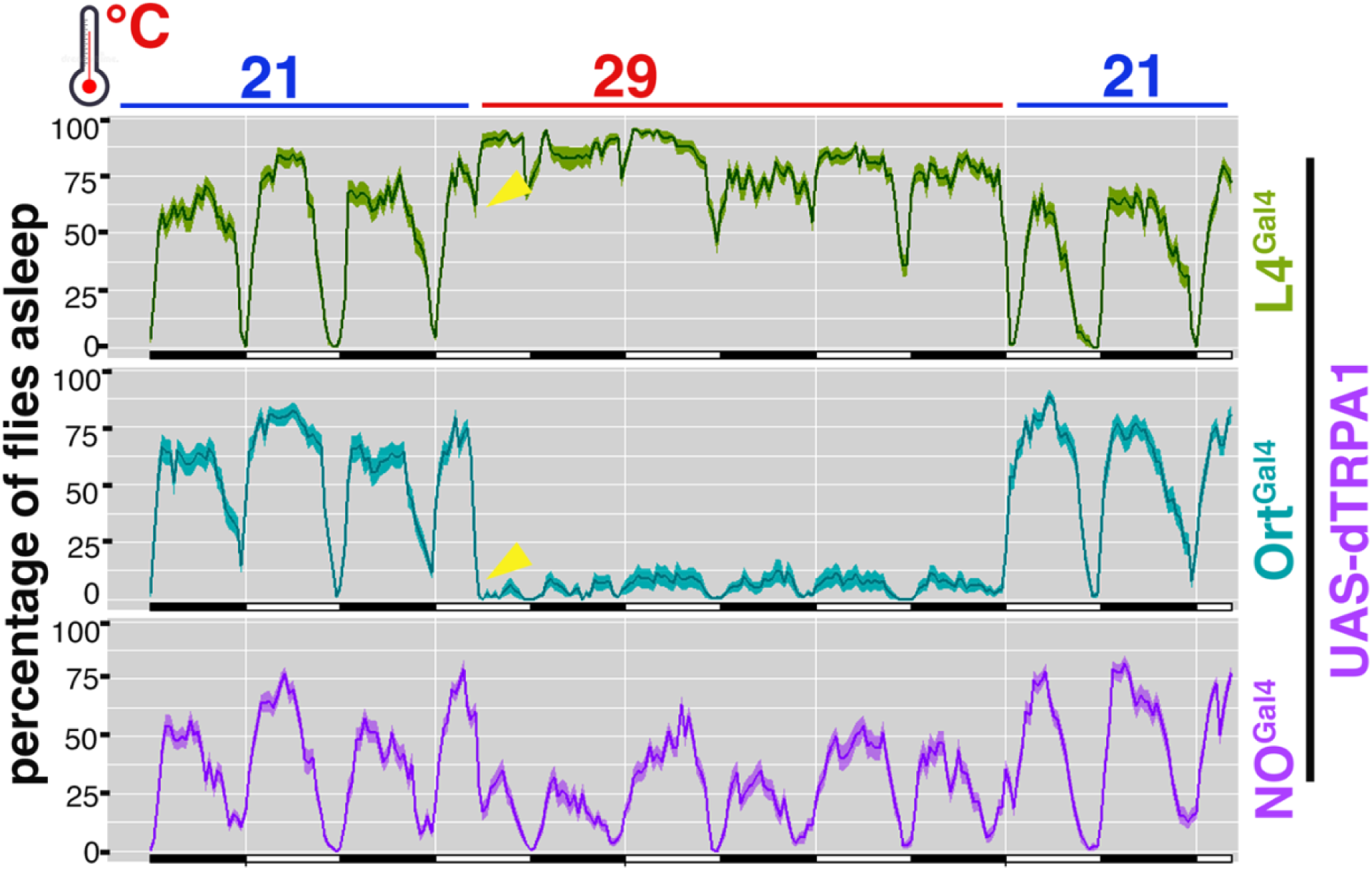
Effect of hyperactivation of lamina neuron on fly sleep behavior. Ethoscope recordings of sleep profile of flies expressing UAS-dTRPA1 driven by L4-Gal4 (green) or ort-GAL4 (Cyan) or without GAL4 (purple), n= 40 per genotype. Restrictive and permissive temperatures are shown on the top, arrow heads indicate the sharp effect on sleep behavior just after switching the temperature. Data shown are combined from two independent experiments with 20 flies each for each genotype. Genotypes are listed in Supplementary Table 1.

**Suppl Fig. 3.**
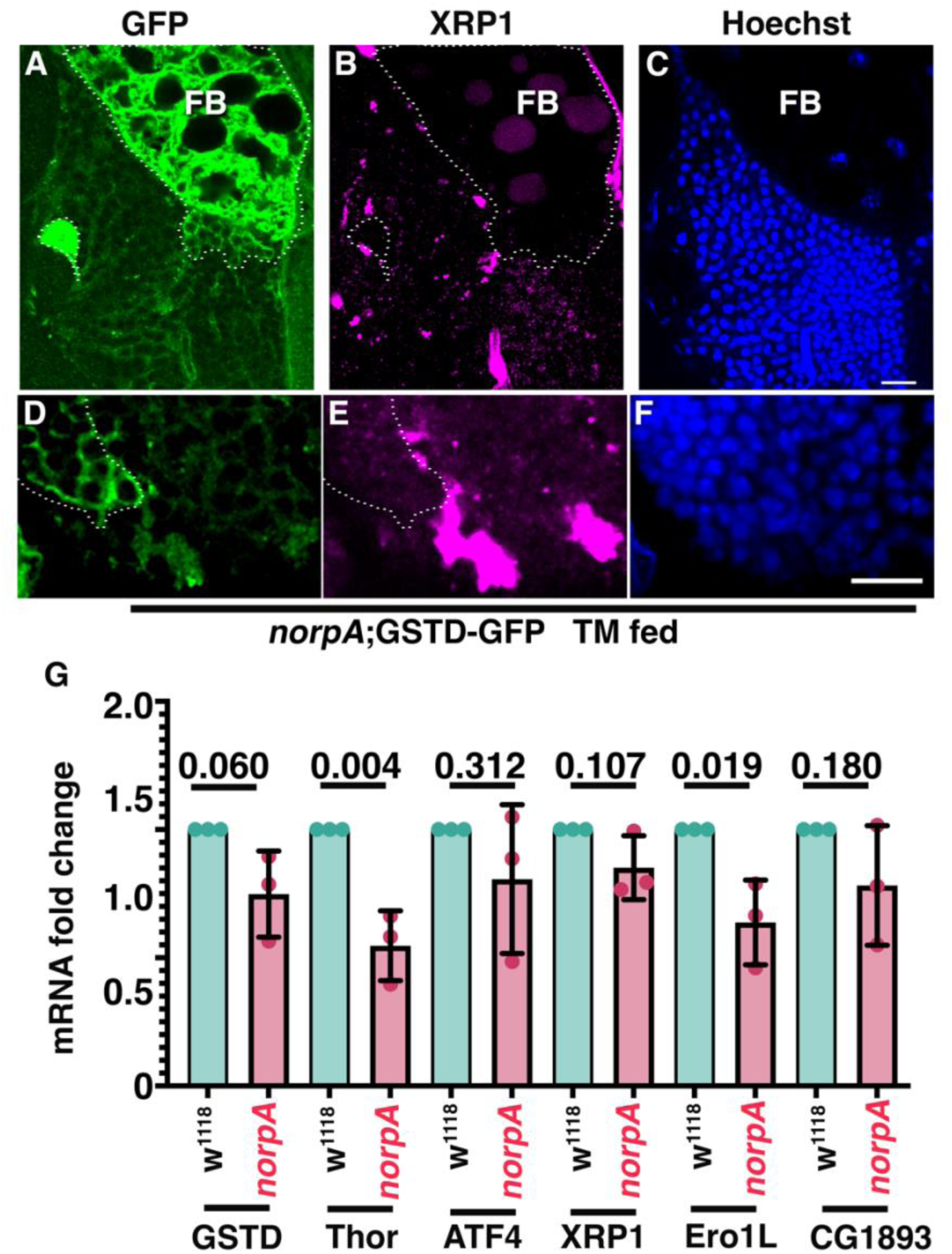
Sequestration of ATF4 and XRP1 in stress granules dampens their responsiveness to stress. (A-F) Adult brain cryosections of TM-fed *norpA* flies expressing GSTD-GFP, a downstream transcriptional target of XRP1, show elevated expression of GSTD-GFP that is limited to cells that lack XRP1-positive SGs (delineated by dotted line). Sections were stained for GFP (A, D), XRP1 (B, E), DNA (C, F). Note the high level of GSTD-GFP expression in fat body cells (FB). Scale bars in C and F are 10 µm. (G) Bar graphs indicate relative mRNA levels for the indicated ISR-related genes in *norpA* and *w*^1118^ control flies as determined by RTqPCR. Expression levels were normalized to Act5c RNA for each genotype and to the levels in *w*^1118^ control flies for each gene. Genotypes are listed in Supplementary Table 1.

**Suppl Fig. 4.**
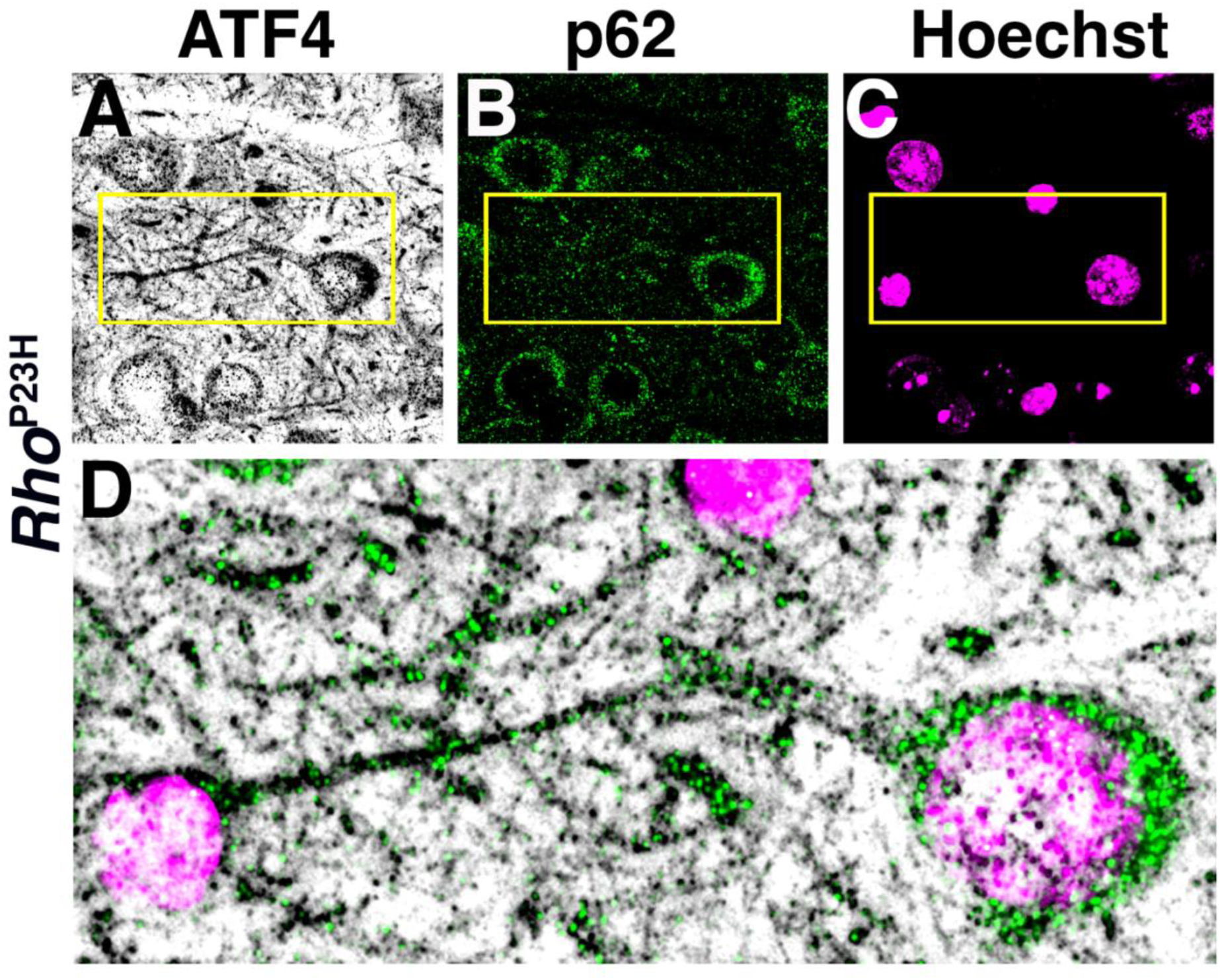
Cytosolic puncta of ATF4 and p62 in neurons of *Rho*^P23H^ mice. (A-D) Sections of thalamus from a 10-week-old *Rho*^P23H^ mouse stained for ATF4 (A), p62 (B) and DNA (C). (D) Merged, magnified image of indicated area from A-C shows ATF4 and p62 punctae at high levels in neurites and cell bodies. Genotype is listed in Supplementary Table 1.

**Supplementary Table 1.**
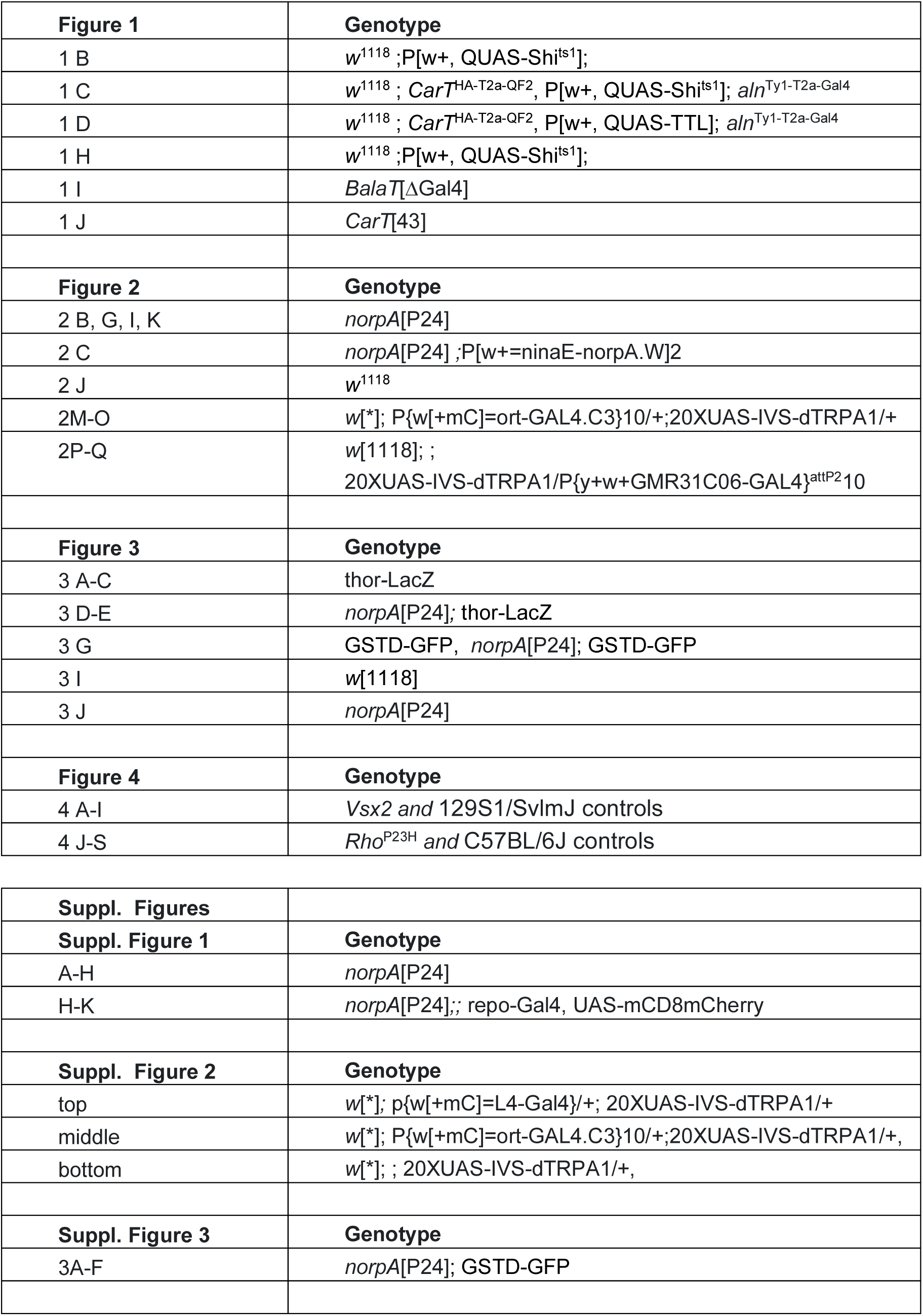
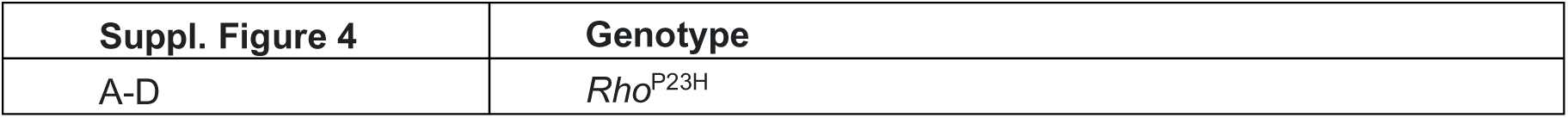
Genotypes of flies or mice used for each figure.

## References

1. G. Livingston et al., Dementia prevention, intervention, and care: 2020 report of the Lancet Commission. Lancet 396, 413–446 (2020).

2. J. R. Ehrlich, J. Goldstein, B. K. Swenor, H. Whitson, K. M. Langa, P. Veliz, Addition of Vision Impairment to a Life-Course Model of Potentially Modifiable Dementia Risk Factors in the US. JAMA Neurol 79, 623–626 (2022).

3. C. Hetz, S. Saxena, ER stress and the unfolded protein response in neurodegeneration. Nat Rev Neurol 13, 477–491 (2017).

4. C. Mathieu, R. V. Pappu, J. P. Taylor, Beyond aggregation: Pathological phase transitions in neurodegenerative disease. Science 370, 56–60 (2020).

5. D. M. Wilson, 3rd, M. R. Cookson, L. Van Den Bosch, H. Zetterberg, D. M. Holtzman, I. Dewachter, Hallmarks of neurodegenerative diseases. Cell 186, 693–714 (2023).

6. J. Labbadia, R. I. Morimoto, The biology of proteostasis in aging and disease. Annu Rev Biochem 84, 435–464 (2015).

7. L. C. Walker, Proteopathic Strains and the Heterogeneity of Neurodegenerative Diseases. Annu Rev Genet 50, 329–346 (2016).

8. W. E. Balch, R. I. Morimoto, A. Dillin, J. W. Kelly, Adapting proteostasis for disease intervention. Science 319, 916–919 (2008).

9. K. Pakos-Zebrucka, I. Koryga, K. Mnich, M. Ljujic, A. Samali, A. M. Gorman, The integrated stress response. EMBO Rep 17, 1374–1395 (2016).

10. S. L. Giandomenico, B. Alvarez-Castelao, E. M. Schuman, Proteostatic regulation in neuronal compartments. Trends Neurosci 45, 41–52 (2022).

11. E. Nachman, P. Verstreken, Synaptic proteostasis in Parkinson’s disease. Curr Opin Neurobiol 72, 72–79 (2022).

12. E. Gomes, J. Shorter, The molecular language of membraneless organelles. J Biol Chem 294, 7115–7127 (2019).

13. B. Wolozin, P. Ivanov, Stress granules and neurodegeneration. Nat Rev Neurosci 20, 649–666 (2019).

14. D. Fujikawa et al., Stress granule formation inhibits stress-induced apoptosis by selectively sequestering executioner caspases. Curr Biol 33, 1967–1981 e1968 (2023).

15. J. R. Buchan, mRNP granules. Assembly, function, and connections with disease. RNA Biol 11, 1019–1030 (2014).

16. Y. Gwon, B. A. Maxwell, R. M. Kolaitis, P. Zhang, H. J. Kim, J. P. Taylor, Ubiquitination of G3BP1 mediates stress granule disassembly in a context-specific manner. Science 372, eabf6548 (2021).

17. S. Shekhar et al., Allnighter pseudokinase-mediated feedback links proteostasis and sleep in Drosophila. Nat Commun 14, 2932 (2023).

18. B. Brown, S. Mitra, F. D. Roach, D. Vasudevan, H. D. Ryoo, The transcription factor Xrp1 is required for PERK-mediated antioxidant gene induction in Drosophila. Elife 10, (2021).

19. T. H. Clausen et al., p62/SQSTM1 and ALFY interact to facilitate the formation of p62 bodies/ALIS and their degradation by autophagy. Autophagy 6, 330–344 (2010).

20. Y. Xu, F. An, J. A. Borycz, J. Borycz, I. A. Meinertzhagen, T. Wang, Histamine Recycling Is Mediated by CarT, a Carcinine Transporter in Drosophila Photoreceptors. PLoS genetics 11, e1005764 (2015).

21. R. Chaturvedi, Z. Luan, P. Guo, H. S. Li, Drosophila Vision Depends on Carcinine Uptake by an Organic Cation Transporter. Cell Rep 14, 2076–2083 (2016).

22. D. Stenesen, A. T. Moehlman, H. Krämer, The carcinine transporter CarT is required in Drosophila photoreceptor neurons to sustain histamine recycling. Elife 4, e10972 (2015).

23. Y. Han, L. Xiong, Y. Xu, T. Tian, T. Wang, The β-alanine transporter BalaT is required for visual neurotransmission in Drosophila. eLife 6, e29146 (2017).

24. B. T. Bloomquist et al., Isolation of a putative phospholipase C gene of Drosophila, norpA, and its role in phototransduction. Cell 54, 723–733 (1988).

25. C. Montell, Drosophila visual transduction. Trends Neurosci 35, 356–363 (2012).

26. C. Gareau et al., Characterization of fragile X mental retardation protein recruitment and dynamics in Drosophila stress granules. PLoS One 8, e55342 (2013).

27. K. Buddika, I. S. Ariyapala, M. A. Hazuga, D. Riffert, N. S. Sokol, Canonical nucleators are dispensable for stress granule assembly in Drosophila intestinal progenitors. J Cell Sci 133, (2020).

28. P. Yang et al., G3BP1 Is a Tunable Switch that Triggers Phase Separation to Assemble Stress Granules. Cell 181, 325–345.e328 (2020).

29. J. Guillén-Boixet et al., RNA-Induced Conformational Switching and Clustering of G3BP Drive Stress Granule Assembly by Condensation. Cell 181, 346–361.e317 (2020).

30. J. D. Ni, L. S. Baik, T. C. Holmes, C. Montell, A rhodopsin in the brain functions in circadian photoentrainment in Drosophila. Nature 545, 340–344 (2017).

31. T. A. Currier, M. M. Pang, T. R. Clandinin, Visual processing in the fly, from photoreceptors to behavior. Genetics 224, (2023).

32. M. Rivera-Alba et al., Wiring economy and volume exclusion determine neuronal placement in the Drosophila brain. Curr Biol 21, 2000–2005 (2011).

33. C. Y. Chow, K. J. Kelsey, M. F. Wolfner, A. G. Clark, Candidate genetic modifiers of retinitis pigmentosa identified by exploiting natural variation in Drosophila. Human molecular genetics 25, 651–659 (2016).

34. C. W. Wittmann et al., Tauopathy in Drosophila: neurodegeneration without neurofibrillary tangles. Science 293, 711–714 (2001).

35. J. R. Sanes, S. L. Zipursky, Design principles of insect and vertebrate visual systems. Neuron 66, 15–36 (2010).

36. C. Bone-Larson et al., Partial rescue of the ocular retardation phenotype by genetic modifiers. J Neurobiol 42, 232–247 (2000).

37. M. Burmeister et al., Ocular retardation mouse caused by Chx10 homeobox null allele: impaired retinal progenitor proliferation and bipolar cell differentiation. Nat Genet 12, 376–384 (1996).

38. S. Sakami et al., Probing mechanisms of photoreceptor degeneration in a new mouse model of the common form of autosomal dominant retinitis pigmentosa due to P23H opsin mutations. J Biol Chem 286, 10551–10567 (2011).

39. H. P. Harding et al., Regulated translation initiation controls stress-induced gene expression in mammalian cells. Mol Cell 6, 1099–1108 (2000).

40. N. Ohoka, S. Yoshii, T. Hattori, K. Onozaki, H. Hayashi, TRB3, a novel ER stress-inducible gene, is induced via ATF4-CHOP pathway and is involved in cell death. EMBO J 24, 1243–1255 (2005).

41. J. Han et al., ER-stress-induced transcriptional regulation increases protein synthesis leading to cell death. Nat Cell Biol 15, 481–490 (2013).

42. L. Baillon, F. Germani, C. Rockel, J. Hilchenbach, K. Basler, Xrp1 is a transcription factor required for cell competition-driven elimination of loser cells. Sci Rep 8, 17712 (2018).

43. J. Blanco, J. C. Cooper, N. E. Baker, Roles of C/EBP class bZip proteins in the growth and cell competition of Rp (’Minute’) mutants in Drosophila. Elife 9, (2020).

44. L. A. Perera et al., Infancy-onset diabetes caused by de-regulated AMPylation of the human endoplasmic reticulum chaperone BiP. EMBO Mol Med 15, e16491 (2023).

45. M. Costa-Mattioli, P. Walter, The integrated stress response: From mechanism to disease. Science 368, (2020).

46. T. Sinnige, A. Yu, R. I. Morimoto, Challenging Proteostasis: Role of the Chaperone Network to Control Aggregation-Prone Proteins in Human Disease. Adv Exp Med Biol 1243, 53–68 (2020).

47. G. P. Sykiotis, D. Bohmann, Keap1/Nrf2 signaling regulates oxidative stress tolerance and lifespan in Drosophila. Dev Cell 14, 76–85 (2008).

48. A. Aguilera-Gomez et al., Phospho-Rasputin Stabilization by Sec16 Is Required for Stress Granule Formation upon Amino Acid Starvation. Cell Rep 20, 935–948 (2017).

49. O. Papoulas et al., dFMRP and Caprin, translational regulators of synaptic plasticity, control the cell cycle at the Drosophila mid-blastula transition. Development 137, 4201–4209 (2010).

50. R. J. Hunt, L. Granat, G. S. McElroy, R. Ranganathan, N. S. Chandel, J. M. Bateman, Mitochondrial stress causes neuronal dysfunction via an ATF4-dependent increase in L-2-hydroxyglutarate. J Cell Biol 218, 4007–4016 (2019).

51. H. Kramer, M. Phistry, Mutations in the Drosophila hook gene inhibit endocytosis of the boss transmembrane ligand into multivesicular bodies. J Cell Biol 133, 1205–1215 (1996).

52. K. N. Richter et al., Glyoxal as an alternative fixative to formaldehyde in immunostaining and super-resolution microscopy. EMBO J 37, 139–159 (2018).

53. K. Pircs et al., Advantages and limitations of different p62-based assays for estimating autophagic activity in Drosophila. PLoS One 7, e44214 (2012).

54. N. Nandi, L. K. Tyra, D. Stenesen, H. Kramer, Stress-induced Cdk5 activity enhances cytoprotective basal autophagy in Drosophila melanogaster by phosphorylating acinus at serine(437). Elife 6, (2017).

55. Q. Geissmann, L. Garcia Rodriguez, E. J. Beckwith, A. S. French, A. R. Jamasb, G. F. Gilestro, Ethoscopes: An open platform for high-throughput ethomics. PLoS Biol 15, e2003026 (2017).

56. Q. Geissmann, L. Garcia Rodriguez, E. J. Beckwith, G. F. Gilestro, Rethomics: An R framework to analyse high-throughput behavioural data. PLoS One 14, e0209331 (2019).

